# A dietary intervention following incretin analog treatment restores adipose tissue functions in diet-induced obese mice

**DOI:** 10.1101/2025.01.08.631845

**Authors:** Joke Seuntjens, Jente De Gols, Bethan K Davies, Fien Van Looy, Ingrid Stockmans, Karen Moermans, Geert Carmeliet, Christophe Matthys, Roman Vangoitsenhoven, Bart Van der Schueren, Steve Stegen, Mitsugu Shimobayashi

## Abstract

Obesity affects more than 15% of the world population and is associated with the development of glucose intolerance and type 2 diabetes. In recent years, incretin analogs are prescribed at a high rate for treatment of obesity and diabetes due to their potent effects on lowering bodyweight and improving glucose homeostasis. However, recent studies suggest that many patients do not stay on incretin analog therapy and thereby rapidly regain bodyweight. The non-compliance of patients to incretin analog therapy is not only due to drug shortage but also insufficient knowledge on the long-term effects of the therapy. To address this knowledge gap and provide a long-term therapy strategy for obesity, we examined the effects of incretin analog treatment and withdrawal on adipose tissue functions in diet-induced obese mice. Our transcriptome data suggest that incretin analog treatment restored most of obesity-mediated deregulated gene expression in adipose tissue. However, genes encoding lipogenic enzymes, downregulated by diet-induced obesity, were not restored by incretin analog treatment. Upon therapy withdrawal, mice displayed rapid bodyweight regain, impaired adipose tissue function, and glucose intolerance. In contrast, a dietary intervention following incretin analog therapy withdrawal restored lipogenic gene expression in adipose tissue, maintained glucose homeostasis, increased lean mass and minimized body weight regain. Our findings revealed the effects of incretin analog therapy and therapy withdrawal on adipose tissue and highlight the importance of a dietary intervention following incretin analog therapy, which may contribute to the development of long-term therapy guidelines of incretin analog therapy for patients with obesity.

## INTRODUCTION

Obesity, caused by an increased positive energy balance, is a metabolic disease characterized by an excessive accumulation of adipose tissue. Today, almost 50% of the world population is overweight, and 15% lives with obesity. The World Health Organization predicts that the prevalence of obesity will increase up to 25% within the next 10 years (1). One of the major obesity-related comorbidities is type 2 diabetes, characterized by high levels of blood glucose and insulin due to insulin resistance (2). Dietary interventions along with physical activities are the first treatment option for obesity. However, long-term efficacy of weight loss by dietary interventions remains a challenge for people with obesity due to so called “obesogenic memory”, which compensates for energy deficit and contributes to weight regain (3–5).

In recent years, incretin analogs have gained attention as a treatment for obesity and diabetes (6). Incretins such as glucagon-like peptide 1 (GLP-1) and glucose-dependent insulinotropic polypeptide (GIP) are gut-derived hormones that slow gastric emptying, reduce appetite, and increase insulin secretion (7, 8). The GLP-1 analog semaglutide and GLP-1/GIP dual analog tirzepatide are the most widely used incretin analogs in clinical practice. Semaglutide and tirzepatide cause approximately 15% and 20 % reductions in bodyweight, respectively, and both drugs improve glucose tolerance and insulin sensitivity (9, 10).

Because of the potent anti-obese and anti-diabetic effects, incretin analogs are currently prescribed at a high rate, resulting in shortages in drug supply (6). Incretin analog therapy is expected to be life-long, which causes financial burdens for patients. Moreover, high dose semaglutide and tirzepatide for obesity treatment was only initiated in clinical practice since 2022 and 2023, respectively. Thus, the long-term adverse effects of these drugs remain to be determined, causing psychological discomforts for patients. Due to these logistic, financial and psychological reasons, many patients do not stay on incretin analog therapy (6, 11). It has been demonstrated that withdrawal of incretin analogs results in rapid bodyweight regain and hyperglycemia (12, 13). Currently, there are no guidelines on how to maintain bodyweight after patients reach target bodyweights upon incretin therapy.

Adipose tissue is the primary energy storage organ. Adipose tissue converts surplus carbohydrates (e.g., glucose) into lipids via *de novo* lipogenesis (DNL) (14). Even though the contribution of adipose DNL to body weight is limited (15), adipose DNL is necessary to maintain whole-body insulin sensitivity and glucose homeostasis (16–19). In addition to the energy storage function, adipose tissue plays an endocrine role to control whole-body energy homeostasis. For example, adipose tissue suppresses food intake by secreting the adipokine leptin (20). Since leptin release is proportional to adipose tissue mass, leptin serves as a key hormone that maintains relatively constant body weight (21). Upon obesity, adipose tissue expands by increasing adipocyte size (hypertrophy), which is associated with inflammatory responses. Obesity also causes impaired adipose function such as decreased adipose DNL, which results in whole-body insulin insensitivity and hyperglycemia (16–19). Obesity-associated leptin resistance results in an increased food intake, and adipocytes in turn produce and secrete more leptin, leading to hyperleptinemia (22). Dietary interventions ameliorate adipocyte hypertrophy, restore adipose DNL, and hyperleptinemia (23, 24). In contrast to dietary interventions, the effects of incretin analogs on adipose tissue at the molecular level remain elusive.

In this study, we characterized the effects of semaglutide and tirzepatide treatment and withdrawal on adipose tissue in high fat diet-induced obese mice. Our transcriptome, histological, biochemical analyses show that incretin analog therapy restored most of adipose tissue functions. Consistent with the recent study demonstrating obesogenic transcriptional memory (5), we identified genes that were still deregulated after incretin therapy-mediated weight loss. Interestingly, genes encoding enzymes in DNL were restored by a dietary intervention (feeding a normal diet) but not by semaglutide and tirzepatide. Semaglutide and tirzepatide therapy withdrawal caused a rapid weight regain and glucose intolerance. In contrast, a dietary intervention following incretin analog therapy restored adipose DNL gene expression and maintained glucose homeostasis. Thus, our data suggest that a dietary intervention following incretin analog therapy is an effective treatment option to restore adipose tissue functions and to circumvent drug shortage and chronic incretin analog treatment.

## MATERIAL AND METHODS

### Animal studies

Male C57BL/6jc mice were fed a normal-chow control diet (ND, 10% fat, ResearchDiets Cat#D12450J) or high-fat diet (HFD, 60% fat, ResearchDiets Cat#D12492) for 8 weeks and then either received daily subcutaneous injections with vehicle (40 mM Tris-HCl, pH8), semaglutide (30 nmol/kg, MedChemExpress) or tirzepatide (10 nmol/kg, MedChemExpress) or HFD was switched to a ND (a dietary intervention), whereas the obesity control group received an HFD continuously. For withdrawal experiments, semaglutide or tirzepatide was removed and mice were either maintained on an HFD or switched to a ND (see in Results section). Mice were housed at 21 °C in a conventional facility with a 14 hour light:10h dark cycle with unlimited access to water and food.

### ARRIVE Guidelines for Animal Experiments

The exact number of animals allocated to each group (n number) is indicated in the figure legends. The sample size was chosen according to our previous studies and published reports in which similar experimental procedures were described. No a priori sample size calculation was performed. In total, 2 animals developed liver cancer, and 2 animals had lipoma, and were excluded from the experiments. The animals were randomly fed with a ND or HFD and allocated to treatment groups with similar distribution of bodyweight. A maximum of four animals were housed together in a cage. The investigators were aware of the group allocation during *in vivo* experiments such as glucose tolerance test and insulin tolerance test and during the data analysis. All outcome measures are defined for each figure. Statistical analyses and a measure of variability are described in the figure legends and Statistics section. The status of animals and experimental procedures are described in this section.

### Insulin tolerance test and glucose tolerance test

For the insulin and glucose tolerance tests, mice were fasted for 6 hours and insulin Humalog (i.p. 0.5 U/kg body weight) or glucose (i.p. 1 g/kg body weight) was administered, respectively. Blood glucose was measured with a blood glucose meter (Contour XT).

### Body composition measurement

Body composition was measured by nuclear magnetic resonance imaging (EchoMRI^TM^).

### Serum measurements

Serum leptin and fasting insulin levels were measured using commercially available ELISA kits (CrystalChem) following the manufacturers’ protocol. Serum free fatty acids were measured using Free Fatty Acid Quantitation kit (Sigma). Serum LDL, cholesterol and triglyceride levels were measured using a chemistry analyzer (DxC 700 AU, Beckman Coulter).

### Histology

Subcutaneous white adipose tissue was fixed in 2% paraformaldehyde, embedded in paraffin, and sliced into 4-µm-thick sections. Tissue sections were stained with hematoxylin and eosin and imaged by Zeiss Axioscan 7 (slide scanner). Adipocyte diameters were quantified with FIJI by using the Adiposoft plugin.

### Immunofluorescent staining

For immunofluorescent staining of white adipose tissue, sections were rehydrated and antigen retrieved by boiling sections in antigen retrieval solution (10 mM sodium citrate, 0.5% Tween 20, pH 6.0). Sections were blocked using 5% BSA and 0.1% triton in PBS and then incubated in primary antibody diluted in diluent with background reducing component (Dako) overnight at 4°C. A primary antibody against tyrosine hydroxylase (1:200, ThermoScientific Cat#PA5-85167) was used. After washing, sections were incubated in secondary antibody (1:1000, Alexa Fluor 568, FisherScientific Cat#10463022), diluted in diluent with background reducing component (Dako). Sections were stained with DAPI for 5min and mounted with Prolong Diamond solution (ThermoScientific). Images were obtained using Zeiss AXIO Imager M2 and analyzed with FIJI. For quantification, fluorophore intensity was measured and normalized to the area of the ROI.

### Immunoblots

Tissues were homogenized in a lysis buffer containing 100 mM Tris (Merck) pH7.5, 2 mM EDTA (Sigma-Aldrich), 2 mM EGTA (Sigma-Aldrich), 150 mM NaCl (Sigma-Aldrich), 1% Triton X-100 (Sigma-Aldrich), cOmplete protease inhibitor cocktail (Sigma-Aldrich) and PhosSTOP (Sigma-Aldrich). Protein concentration was determined by the Bradford assay (Bio-Rad), and equal amounts of protein were separated by SDS-PAGE and transferred onto nitrocellulose membranes (Sigma-Aldrich). Antibodies used in this study were as follows: HK2 (1:1000, Cat#2867), FASN (1:1000, Cat#3189), ACC (1:1000, Cat#3662), ACLY (1:1000, Cat#13390), AKT (1:1000, Cat#4685) and AKT-S473 (1:1000, Cat#4060) from Cell Signaling Technology, CALX (1:5000, Cat#PA-34754) from Invitrogen and TH (1:1000, Cat#PA5-85167) from ThermoScientific. A fluorescently labeled secondary goat anti-rabbit antibody (1:20.000, Cat#926-32211) from Licor was used. For quantification, the signals were normalized to a loading control (CALX).

### RNA isolation and quantitative real-time PCR

Total RNA was extracted from the tissues using Nucleozol reagent (Macherey-Nagel) and Nucleospin kit (Macherey-Nagel). cDNA was synthesized from 500 ng RNA using iScript cDNA synthesis kit (BioRad). Gene expression levels were assessed by QuantStudio (Applied Biosystems) using Fast SYBR Green Master Mix (ThermoScientific) according to the manufacturer’s instructions. mRNA levels were normalized to the expression of *Tbp* or *18S*. Primers used for RT-qPCR are listed in Table S1.

### RNA sequencing

Total RNA was extracted from 8 mice for each group, using Nucleozol reagent (Macherey-Nagel) according to the manufacturer’s protocol. The RNA concentration was measured using a NanoDrop™ Spectrophotometer (Thermo Fisher Scientific). RNA integrity was evaluated with an Agilent Bioanalyzer. The RNA samples were processed by the Genomics Core Leuven (Belgium). Libraries were generated with the Illumina TruSeq-Stranded mRNA Sample Preparation Kit and subsequently sequenced on the Illumina HiSeq 4000. Reads of 50 bp were generated, and an average of 4 million assigned reads was obtained. One sample from the ND control group (outlier on PCA plot) and one from the ND semaglutide group (poor read counts) were excluded from the analysis. The reads were mapped against the mouse genome mm10. Differential gene expression analysis was performed with DESeq2 package in RStudio.

### Study Approval

All animal experiments were approved by the the Animal Ethics Committee from KU Leuven (P214/2023).

### Statistics

All data are shown as the means ± SEM. Sample numbers are indicated in each figure legend. To determine the statistical significance between groups, one-way ANOVA and Tukey’s multiple comparison test was performed. All statistical analyses were performed using GraphPad Prism 9 (GraphPad Software). p < 0.05 was considered as statistically significant.

## RESULTS

### Incretin analogs restore most of adipose tissue functions

To study the effect of incretin analogs on adipose tissue function, we first fed mice with an high fat diet (HFD) for 8 weeks and subsequently treated them daily with s.c. semaglutide and tirzepatide for 4 weeks in the presence of an HFD. We compared the efficacy of these drugs to a dietary intervention (diet switch from an HFD to a normal diet (ND)) (**Fig. 1A**). Semaglutide caused 24% bodyweight reduction in HFD-fed obese mice (**Fig. 1B-C**). Tirzepatide resulted in more than 32% weight loss, reaching the bodyweight of ND-fed control mice (**Fig. 1B-C**). Consistent with bodyweight loss, semaglutide and tirzepatide treatment decreased fat mass and adipocyte size (**Fig. 1D-F** and **S1A**). HFD-induced loss of relative lean mass was restored by tirzepatide, and there was a trend towards restored lean mass by semaglutide (**Fig. S1B**). It has been demonstrated that incretin analogs cause a decrease in bodyweight by suppressing appetite and food intake (25). Indeed, incretin analog treatment reduced food and energy intake (**Fig. 1G** and **S1C**). In line with human data (9, 10), fasting insulin levels, insulin sensitivity, and glucose tolerance were restored upon incretin analog treatment, similar to a dietary intervention (**Fig. 1H-I** and **S1D-G**). Furthermore, serum free fatty acids were reduced and the serum lipid profile (low-density lipoprotein (LDL), cholesterol and triglycerides) was improved upon incretin analog treatment (**Fig. S1H-I**). These data confirmed the efficacy of semaglutide and tirzepatide for the treatment of obesity.

**Figure 1.**
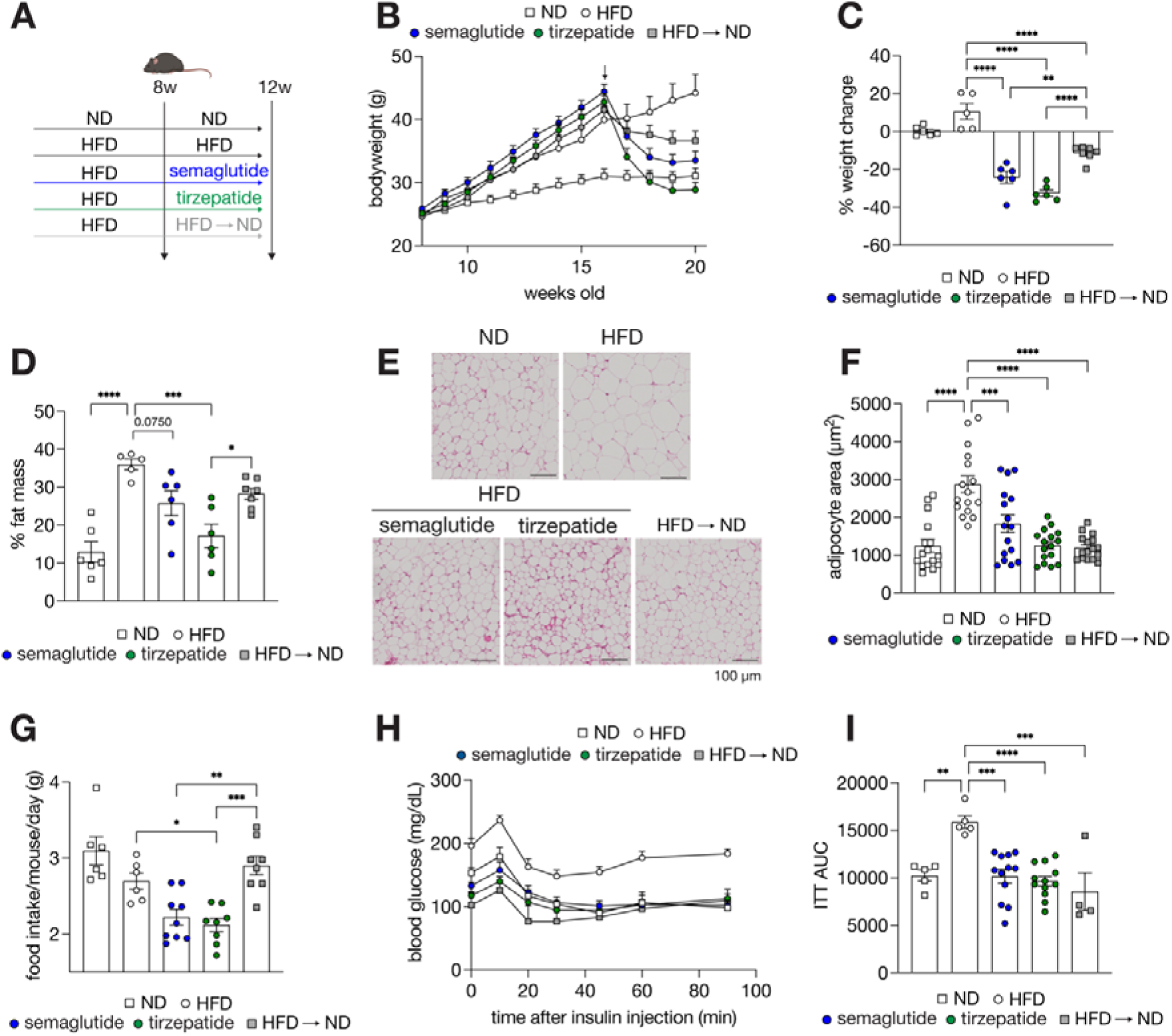
Incretin analog treatment reduces bodyweight and adiposity and restores insulin sensitivity and glucose tolerance. **A.** Experimental design of incretin analog treatment. **B-D**. Bodyweight (B), percentage of weight change (C) and relative fat mass (D) 4 weeks after incretin analog treatment or a dietary intervention. n = 6 (ND, SEM, TZP), 5 (HFD) and 7 (HFD ➔ ND). **E.** Hematoxylin and eosin staining of sWAT. bar = 100 µm. **F**. Quantification of adipocyte area shown in Fig. 1D. Two sections for each animal were analyzed using Adiposoft plugin in ImageJ and the median of adipocyte area was calculated for each slice. n = 8. **G**. Food intake was measured per cage and averaged per mouse after 4 weeks of incretin analog treatment or dietary intervention. N = 6 (ND and HFD), 9 (SEM), and 8 (TZP and HFD ➔ ND). **H-I**. Blood glucose levels during insulin tolerance test (H) and Area under the curve (AUC) (I). 0.5U/kg insulin was injected, and glucose concentrations were measured at the indicated timepoints. n = 5 (ND and HFD), 12 (SEM and TZP) and 4 (HFD ➔ ND). ND: normal diet, HFD: high-fat diet, SEM: semaglutide, TZP: tirzepatide and HFD ➔ ND: dietary intervention. *p<0.05, ** p<0.01, ***p<0.001, ****p <0.0001, One-Way ANOVA.

To study the effect of incretin analog treatment on adipose tissue at the molecular level, we examined the transcriptome of subcutaneous white adipose tissue (sWAT) by bulk RNA sequencing. Our transcriptome data showed that 180 genes and 198 genes (log2 fold (log2FC > |±1|, p<0.05) were upregulated and downregulated in HFD-fed obese mice, respectively (**Fig. 2A-D**). The upregulated genes are known to be involved in PPAR signaling and beta oxidation (*Fabp5*, *Acsl1*, *Angptl4* and *Pltp)* and inflammation (*Cd68, Adgre1* and *Ccl6*) (**Fig. S2A**). As previously described, we also observed increased expression of the gene encoding leptin (*Lep*) (**Fig. 2E**), probably due to obesity-induced leptin resistance (26, 27). These upregulated genes in HFD-fed mice were restored by semaglutide, tirzepatide or a dietary intervention (**Fig. 2E-F** and **S2B**). Consistent with the normalized *Lep* expression, hyperleptinemia caused by an HFD was also restored upon semaglutide, tirzepatide or a dietary intervention (**Fig. S2C**). Pro-inflammatory markers (*Cd68* and *Ccl2*) were also reduced in visceral WAT (vWAT), suggesting that adipose inflammation was resolved upon incretin analog treatment (**Fig. S2D**). HFD-induced downregulated genes are related to fatty acid metabolism and the selenium micronutrient network (**Fig. 2B** and **S2A**). Among the 198 downregulated genes in HFD-fed mice, semaglutide and tirzepatide restored 120 and 145 genes, respectively, including genes from the selenium micronutrient network (*Fads1*, *Gpx3* and *Kmo*) (**Fig. 2B, 2D,** and **2G**). Interestingly, we also observed that tirzepatide specifically rescued some of the downregulated genes. These genes are involved in fatty acid desaturation (*Scd3*, *Scd4* and *Fads1*), glycogen metabolism (*Phkg1*, *Pygm* and *Pygl*), glycolysis (*Pfkm* and *Eno3*), and sympathetic activity (*Adrb3*) (**Fig. S2E**). This tirzepatide-specific rescue might be explained by expression of the GIP receptor in adipose tissue (28). HFD-induced obesity is known to cause sympathetic denervation in adipose tissue (29). Thus, we also examined sympathetic innervation in adipose tissue by examining the expression of the catecholamine synthesis enzyme tyrosine hydroxylase (TH), commonly used as a sympathetic marker (29, 30). TH expression was reduced upon an HFD and restored by semaglutide and tirzepatide (**Fig. 2H-I** and **S2F-G**). Overall, these data suggest that incretin analogs, especially tirzepatide, restore most of the obesity-mediated impaired adipose tissue function.

**Figure 2.**
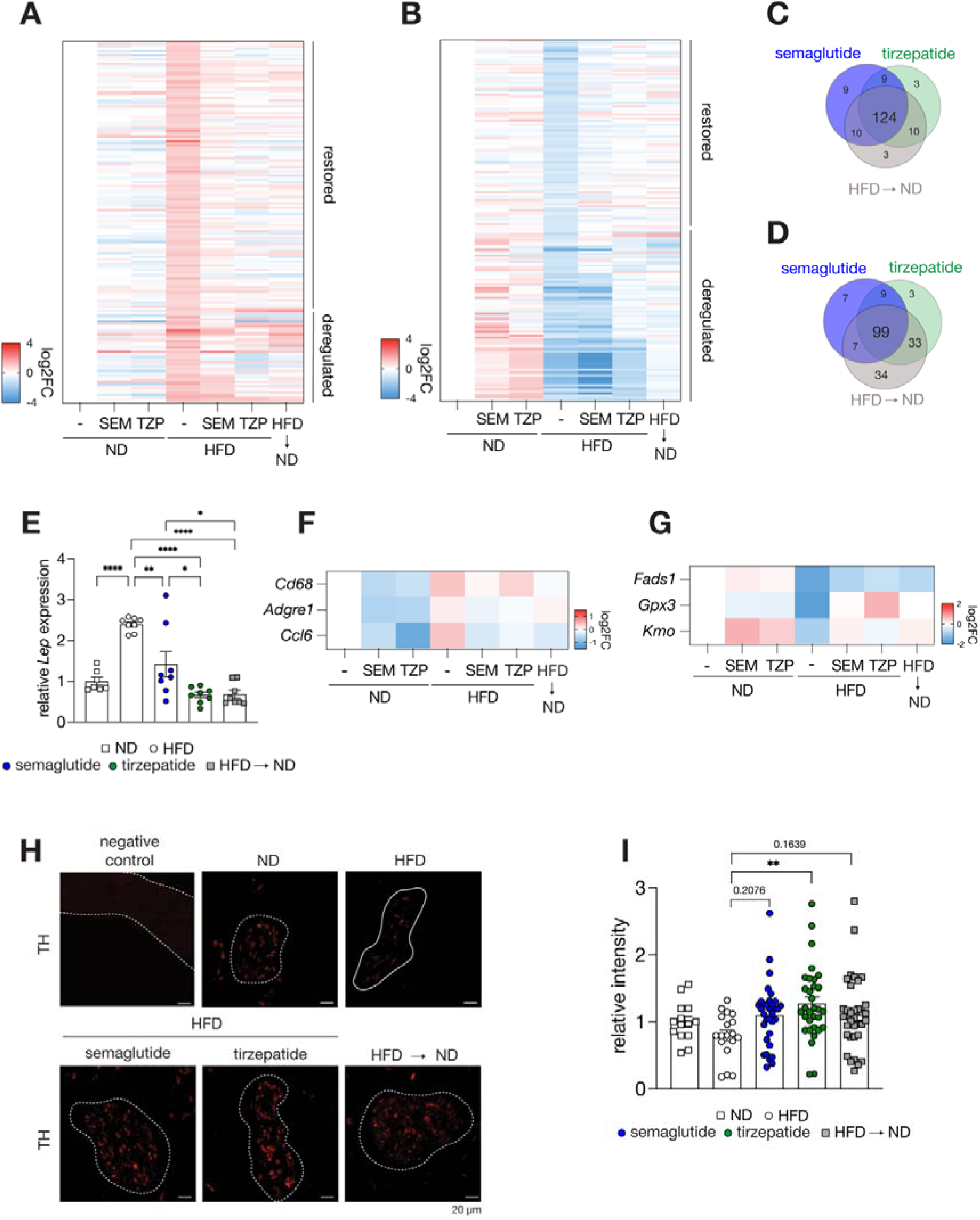
Incretin analogs restore most of the HFD-induced deregulated functions in adipose tissue. **A-B**. Relative mRNA expression of genes that were upregulated (A) or downregulated (B) upon diet-induced obesity and the effects of incretin therapy or a dietary intervention. n = 7 (ND, and ND SEM), n = 8 (ND TZP, HFD, HFD SEM, HFD TZP, and HFD ➔ ND). **C-D**. Venndiagrams showing the number of genes that was upregulated (C) and downregulated (D) upon diet-induced obesity and restored by incretin therapy or dietary interventions. **E-G**. Relative expression of *Lep* mRNA (encoding leptin) (E), pro-inflammatory markers (F), and genes involved in selenium micronutrient pathway (G). n = 7 (ND, and ND SEM), n = 8 (ND TZP, HFD, HFD SEM, HFD TZP, and HFD ➔ ND). **H-I**. Immunostaining of TH-expressing sympathetic nerves on sWAT. Nerve bundles are indicated with a dotted line (H) and quantification (I). N = 14 (ND), N = 18 (HFD), N = 33 (semaglutide), N = 33 (tirzepatide), N = 35 (HFD ➔ ND) *p<0.05,** p<0.01, ****p<0.0001, One-Way ANOVA.

### Deregulated transcriptional signature that is retained upon weight loss

It has been recently shown that adipose tissue possesses an epigenetic memory, which explains the deregulation of some transcripts even after significant weight loss by dietary intervention or bariatric surgery (5). Among the 378 deregulated genes, 104 genes, 74 genes and 51 genes remained deregulated after weight loss with semaglutide, tirzepatide or a dietary intervention, respectively (**Fig. 3A** and **2A-B**). Our transcriptome data also confirmed the retained deregulation of obesogenic-marker genes identified in (5), upon incretin analog treatment (**Fig. 3B**). *Ctsc*, *Maob* and *Dnah3* were identified to be downregulated and *Ctsd*, *Apobec1*, *Mmp12* and *Gpnmb* were identified to be upregulated in the adipocyte fraction of adipose tissue after weight loss (5). Notably, some of the downregulated marker genes were only restored by a dietary intervention and not by incretin analog treatment (**Fig. 3B**). These data confirm the presence of an obesogenic transcriptional memory as described in the recent study (5) and further suggest that this memory may be influenced by the type of prior obesity treatment.

**Figure 3.**
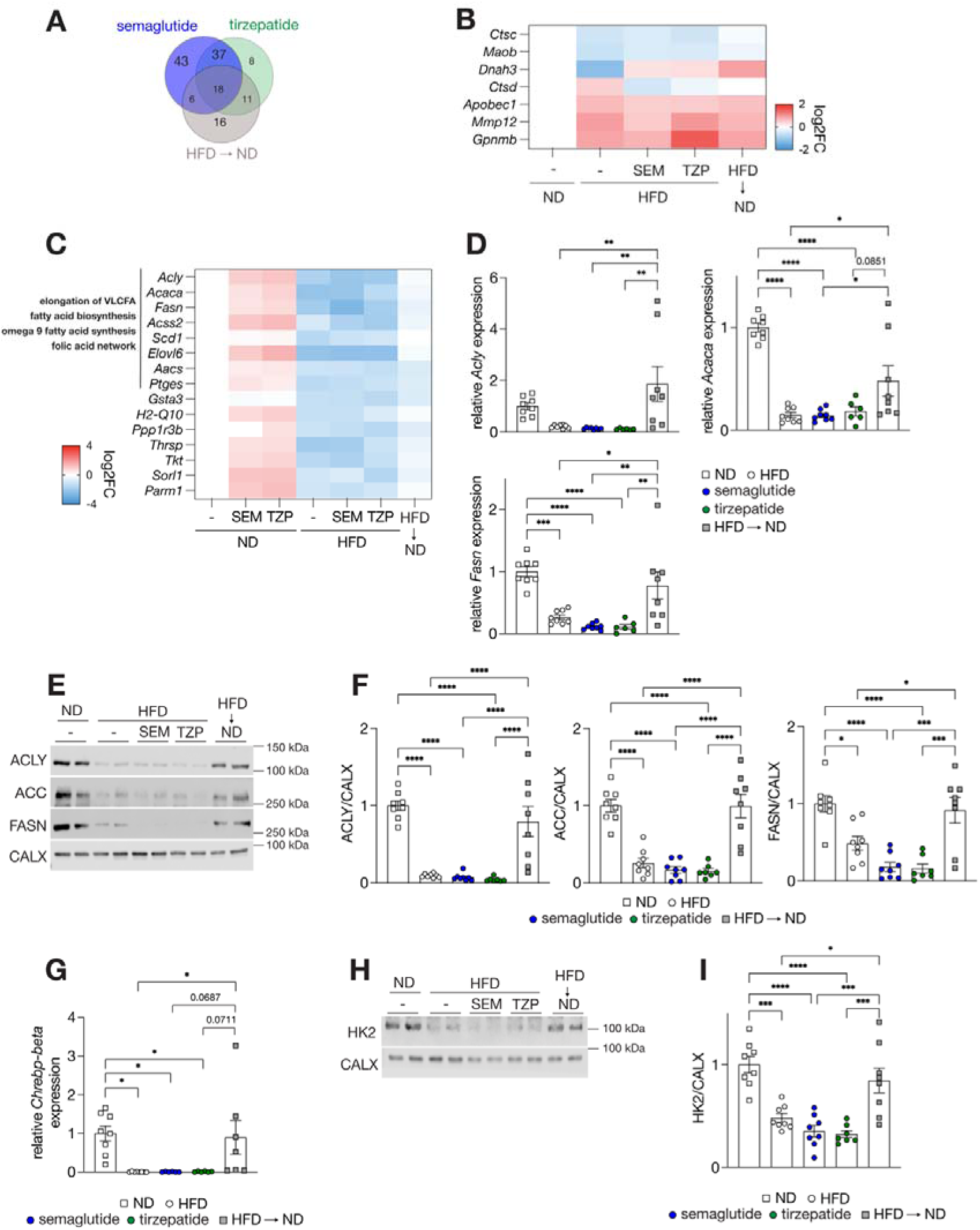
Deregulated transcriptional signature remains after incretin analog therapy. **A.** The number of genes that are deregulated upon an HFD and remained deregulated upon semaglutide, tirzepatide or a dietary intervention. **B**. Relative expression of genes involved in obesogenic memory (5). n = 7 (ND, and ND SEM), n = 8 (ND TZP, HFD, HFD SEM, HFD TZP, and HFD ➔ ND). **C**. Relative expresssion of genes specifically restored by a dietary intervention. n = 7 (ND, and ND SEM), n = 8 (ND TZP, HFD, HFD SEM, HFD TZP, and HFD ➔ ND). **D**. Relative expression of genes encoding ATP citrate lyase (*Acly*), acetyl-CoA carboxylase (*Acaca*), and fatty acid synthase (*Fasn*) in sWAT. n = 8. **E-F**. Immunoblot analysis of lipogenic enzymes in sWAT 4 weeks after incretin analog treatment or dietary intervention (E) and quantification (F). CALX serves as a loading control. For quantification, intensities were normalized to CALX. n = 8. **G**. Relative expression of carbohydrate response element binding protein-beta isoform (*Chrebp*-*beta*). n =7 (HFD ➔ ND) and 8 (ND, HFD, SEM and TZP). **H-I**. Immunoblot analysis of hexokinase 2 (HK2) (H) and quantification (I) n = 8. sWAT: subcutaneous white adipose tissue. *p<0.05,** p<0.01,***p<0.001, ****p<0.0001, One-Way ANOVA.

### Incretin analog treatment does not restore the expression of DNL genes in adipose tissue

In our transcriptome data of sWAT, we noted that the decreased expression of genes encoding enzymes in fatty acid biosynthesis (*Acly*, *Acss2*, *Fasn*, *Acacb* and *Acaca*) and elongation of very long chain fatty acids (*Scd1*, *Scd3*, *Scd4*, *Elovl6* and *Fads1*) was restored by a dietary intervention, but not by incretin analogs (**Fig. 3C**). We confirmed that mRNA expression of *Acly*, *Acaca* and *Fasn* and their protein levels were restored by a dietary intervention, but not by incretin analogs (**Fig. 3D-F**). A similar effect was observed in vWAT and brown adipose tissue (**Fig. S3A-F**). The lack of rescue for the DNL genes is not due to short-term treatment of incretin analogs since longer semaglutide or tirzepatide treatment (8 weeks) also did not restore expression of DNL genes and proteins (**Fig. S3G-I**). These data suggest that DNL was not restored in adipose tissue upon incretin analog therapy. In contrast to HFD-fed obese mice, incretin analog treatment in ND-fed mice caused increased expression of these DNL genes and their encoding proteins (**Fig. S4A-C**). Thus, incretin analogs have beneficial effects on fatty acid metabolism in ND-fed mice, but these beneficial effects are abolished in an HFD-fed condition.

Next, we investigated the underlying mechanism of unrestored DNL gene expression. Insulin signaling and glucose influx are known to promote expression of DNL genes. As previously reported (19, 31), insulin signaling was not affected by an HFD, incretin analogs nor a dietary intervention, as examined by AKT phosphorylation in sWAT (**Fig. S3J-K**). The transcriptional factor Carbohydrate Responsive Element Binding Protein (ChREBP) promotes the expression of DNL genes in response to glucose (32). Expression of *Chrebp-beta,* an isoform shown to be downregulated upon HFD and an important contributor to insulin sensitivity (17), was not restored upon incretin analog treatment (**Fig. 3G**). It has been shown that adipose HK2 expression is decreased upon diet-induced obesity, which results in reduced expression of *Chrebp-beta* and its target DNL genes (18). We found that HK2 protein expression was restored by a dietary intervention, but not by incretin analog treatment (**Fig. 3H-I**). These data suggest that incretin analog therapy does not restore the expression of genes involved in adipose DNL, most likely due to low glucose influx caused by reduced HK2 expression and thereby reduced ChREBP activity.

### Incretin analog therapy withdrawal impairs adipose tissue function

We noted that the weight loss effects of semaglutide and tirzepatide reached a plateau after 3 weeks of treatment (**Fig. 1A** and **4B**). Thus, we next investigated the effect of incretin therapy withdrawal. HFD-fed obese mice were treated with semaglutide or tirzepatide for 4 weeks, followed by therapy withdrawal (**Fig. 4A** and **S5A**). Tirzepatide withdrawal resulted in 60% bodyweight regain 4 weeks after therapy withdrawal, which was significantly higher than observed in mice after withdrawal from semaglutide or a dietary intervention (**Fig. 4B-C** and **S5B-C**). In all therapy withdrawal conditions, mice rapidly regained their bodyweight in the first week, most likely to compensate for the ‘energy deficit’ caused by incretin analog treatment or a dietary intervention (**Fig. 4D** and **S5D**). The regained bodyweight was due to an increase in fat mass, adiposity, and adipocyte size (**Fig. 4E** and **S5E-H**). Regardless of their prior therapy, therapy-withdrawn mice became glucose intolerant and insulin resistant as shown by hyperinsulinemia (**Fig. 4F-H** and **S6A-C**). We further examined the effects of incretin therapy withdrawal on adipose tissue function. We observed increased expression of inflammatory markers (*Ccl2* and *Cd68*) in vWAT upon incretin analog withdrawal (**Fig. 4I** and **S6D-E**). Also, incretin analog withdrawal caused an increase in *Lep* mRNA levels, which was associated with hyperleptinemia (**Fig. 4J-K** and **S6F-G**). Food intake was increased 4 weeks after incretin analog therapy withdrawal (**Fig. S6H**). The serum lipid profiles (cholesterol, LDL and triglycerides) were worsened by therapy withdrawal (**Fig. S6I**). Overall, our data suggest that incretin therapy withdrawal rapidly impairs adipose tissue function and whole-body glucose homeostasis. Thus, our findings highlight the requirement of a follow-up therapy after incretin therapy.

**Figure 4.**
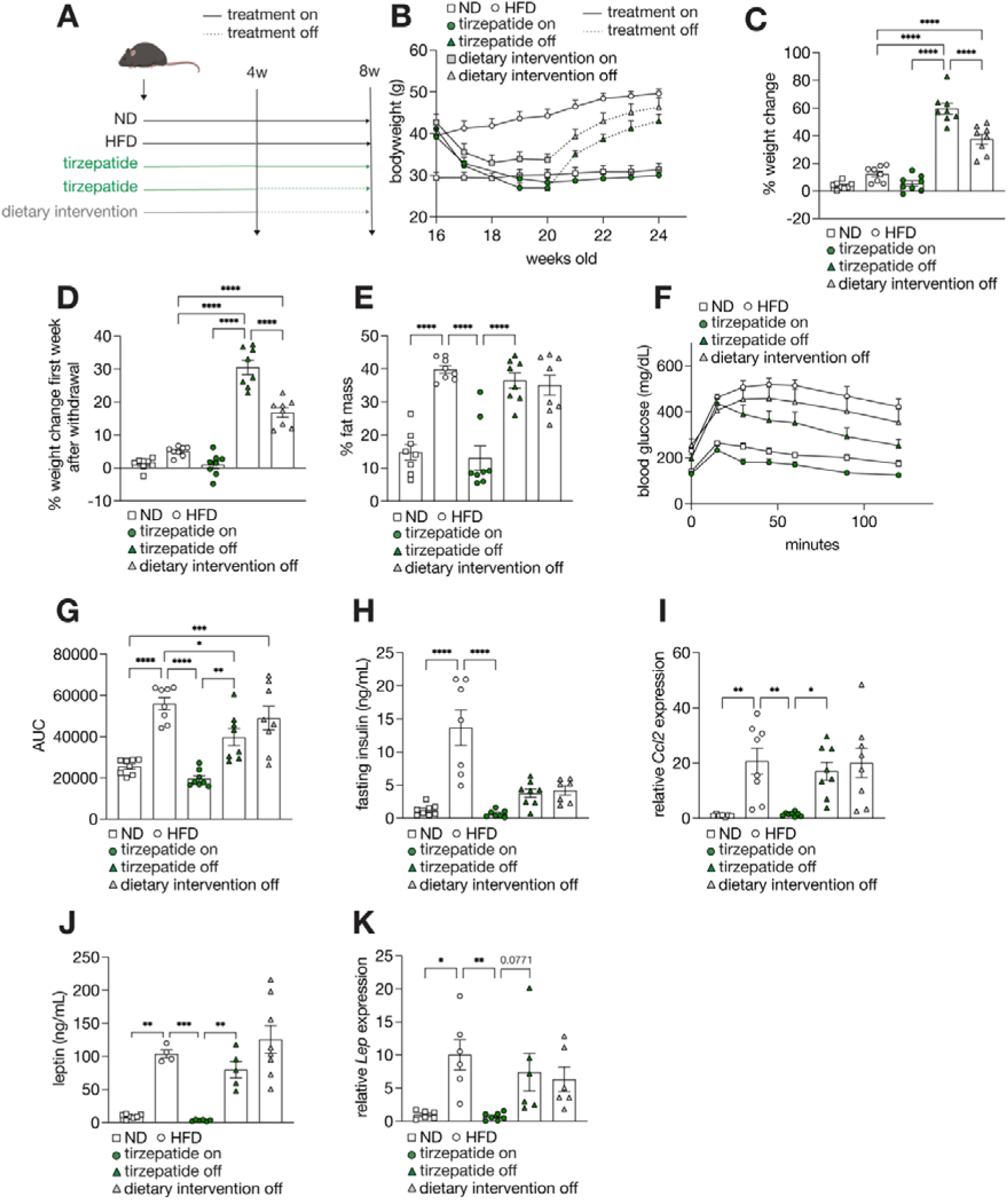
Tirzepatide withdrawal impairs adipose tissue function. **A.** Experimental design of therapy withdrawal. **B**. Bodyweight curve during continuous TZP treatment or before and after therapy withdrawal with diet switch to HFD as control or continuous TZP treatment. n = 8. The dotted line indicates therapy withdrawal. **C-D**. Percentage of bodyweight change 4 weeks after therapy withdrawal compared to start of withdrawal (C) and after the first week of therapy withdrawal (D). n = 8. **E**. Relative fat mass after therapy withdrawal or continuous TZP treatment. n = 8. **F-G**. Blood glucose levels during glucose tolerance test after 4 weeks afterof therapy withdrawal or continuous TZP treatment (F) and AUC (G). 1g/kg glucose was injected, and glucose concentrations were measured at the indicated timepoints. n = 8. **H**. Fasting insulin (ng/mL) levels 4 weeks after therapy withdrawal or continuous TZP treatment. n = 7 (HFD, TZP off, dietary intervention off) and 8 (ND, TZP). **I**. Relative expression of the pro-inflammatory marker *Ccl2* in vWAT. n = 8. **J**. Serum leptin levels (ng/mL). n = 8 (ND, TZP, dietary intervention off), 4 (HFD) and 5 (TZP off). **K**. Relative expression of *Lep* (encoding leptin) in sWAT 4 weeks after therapy withdrawal or continuous TZP treatment. n = 6 (ND, HFD, TZP off, dietary intervention off) and 7 (TZP). *p<0.05,**p<0.01, ***p<0.001, ****p<0.0001, One-Way ANOVA.

### A dietary intervention after incretin analog treatment withdrawal restores adipose tissue functions and maintains glucose homeostasis

In clinical practice, patients are advised to follow energy-restricted diets during and after incretin analog treatment (33). As far as we are aware of, there is no study on the effects of a dietary intervention after incretin therapy withdrawal in either human or mice. Thus, we investigated the effects of a dietary intervention in mice that were withdrawn from incretin analog therapy. HFD-fed mice were first treated with semaglutide and tirzepatide for 4 weeks. Half of the mice were maintained on an HFD, and the other half was switched to a ND (a dietary intervention) upon incretin therapy withdrawal (**Fig. 5A** and **S7A**). In contrast to mice maintained on HFD, mice switched to ND gained significantly less bodyweight (**Fig. 5B-C** and **S7B-C**). Most of the bodyweight regain was observed during the first week after incretin analog withdrawal (**Fig. S7D**), which correlates with the increased food intake one week after therapy withdrawal (**Fig. 5D** and **Fig. S7E**). The change in bodyweight was reflected by a change in total fat mass and adipose depot weight (**Fig. 5E** and **S7F-H**). Notably, after tirzepatide withdrawal, mice receiving a ND still gained almost 14% of their bodyweight 4 weeks after therapy withdrawal (**Fig. 5C**), which correlates with a significant increase in lean mass (**Fig. 5F** and **S7I-J**). A dietary intervention following incretin therapy withdrawal, maintained glucose tolerance and insulin sensitivity (**Fig. 5G-H** and **S8A-F**). Compared to mice fed with HFD following incretin analog withdrawal, ND-fed mice after incretin analog withdrawal displayed a low expression of the inflammatory marker *Ccl2* and normal serum leptin levels (**Fig. 5I** and **S8G-I**). Furthermore, in contrast to HFD, ND feeding following incretin therapy withdrawal restored the expression of DNL genes and proteins in adipose tissue (**Fig. 5J-K** and **S9A-D**). These data highlight the importance of a dietary intervention after incretin analog therapy withdrawal to fully restore adipose tissue function, minimize bodyweight regain, and maintain glucose tolerance.

**Figure 5.**
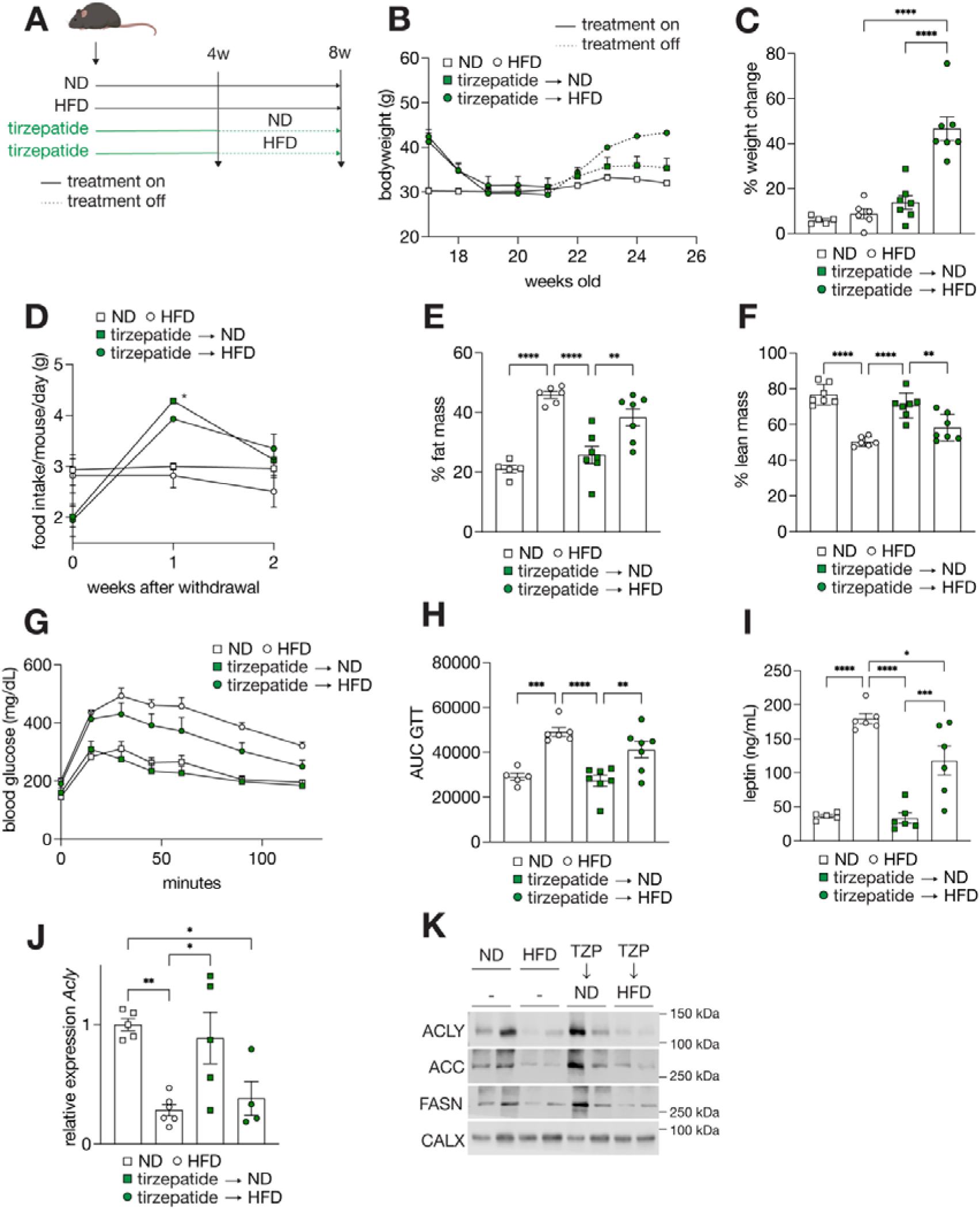
A dietary intervention following tirzepatide treatment restores adipose tissue function and maintained glucose tolerance. **A.** Experimental design of dietary intervention after incretin withdrawal. **B**. Bodyweight of mice that were maintained on a HFD or switched to a ND for 4 weeks after TZP withdrawal. The dotted line indicates therapy withdrawal. n = 5 (ND), 6 (HFD) and 7 (TZP ➔ ND, TZP ➔ HFD). **C.** Percentage of bodyweight change 4 weeks after TZP withdrawal. n = 5 (ND), 6 (HFD) and 7 (TZP ➔ ND, TZP ➔ HFD). **D**. Food intake in the first and second week after TZP withdrawal. **E-F**. Relative fat mass (E) and lean mass (F) 4 weeks after tirzepatide withdrawal. n = 5 (ND), 6 (HFD) and 7 (TZP ➔ ND, TZP ➔ HFD). **G-H**. Blood glucose levels during glucose tolerance test performed in mice 4 weeks after TZP withdrawal (G) and AUC (H). 1g/kg glucose was injected, and glucose concentrations were measured at the indicated timepoints. n = 5 (ND), 6 (HFD) and 7 (TZP ➔ ND, TZP ➔ HFD). **I**. Serum leptin 4 weeks after TZP withdrawal. n = 5 (ND), 6 (HFD, TZP ➔ ND, TZP ➔ HFD). **J**. Relative expression of *Acly* after TZP withdrawal. n = 5 (ND), 6 (HFD) and 7 (TZP ➔ ND, TZP ➔ HFD). **K**. Immunoblot analysis of ACLY, ACC, FASN in sWAT 4 weeks after TZP withdrawal. CALX serves as a loading control. n = 5 (ND), 6 (HFD, TZP ➔ ND, TZP ➔ HFD). *p<0.05,**p<0.01, ***p<0.001, ****p<0.0001, One-Way Anova

## DISCUSSION

In this study, we investigated the effects of the incretin analogs semaglutide and tirzepatide on adipose tissue function in HFD-fed obese mice. We show that incretin analogs restored most of the adipose tissue functions, but not adipose DNL gene expression. Incretin therapy withdrawal resulted in rapid body weight regain, impaired adipose tissue functions, and glucose intolerance. In contrast, a dietary intervention following incretin analog therapy restored adipose DNL expression, minimized bodyweight gain, and maintained glucose tolerance. Our findings suggest that incretin therapy to treat people with obesity needs to be chronic or followed by a dietary intervention.

It was recently shown in humans and mice that adipocytes retain epigenetic memory of obesity even after significant weight loss (5). According to this recent study, the obesogenic memory in adipose tissue contributes to the rapid bodyweight regain after dietary interventions. Our study also identified obesity-mediated deregulated genes that were not restored after incretin therapy- or a dietary intervention-mediated weight loss (**Fig. 2A-B** and **3A**). Similar to a dietary intervention, incretin analog therapy withdrawal caused a rapid bodyweight regain and development of glucose intolerance. The rate of bodyweight regain upon therapy withdrawal was faster than the initial bodyweight gain by an HFD (**Fig. 4D**). These data support the presence of an obesogenic memory in adipose tissue (5). Interestingly, our transcriptome data in adipose tissue also identified HFD-mediated deregulated genes that were only restored by a dietary intervention and not by semaglutide or/and tirzepatide. It needs to be determined whether a therapy-specific epigenetic memory explains the unresponsiveness of these genes to incretin analog therapy.

The expression of DNL genes and proteins was restored by a dietary intervention but not after incretin therapy-mediated weight loss, most likely because HK2 expression and ChREBP activity remained downregulated in semaglutide- or tirzepatide-treated mice (**Fig. 3G-I**). The decreased HK2 expression and ChREBP activity might reflect an incretin therapy-specific obesogenic memory. Alternatively, the presence of a lipid rich and calorie-dense HFD might contribute to the decreased expression of HK2 and subsequently DNL genes. Since we observed restored HK2 expression, ChREBP activity, and DNL gene expression upon a dietary intervention, incretin analog therapy combined with a dietary intervention might have more potent anti-obese and anti-diabetic effects.

Adipose DNL has been shown to promote whole-body insulin sensitivity and glucose tolerance by directly contributing to glucose clearance and indirectly suppressing glucose production in the liver (16–19). Our data demonstrate that adipose DNL gene expression was not restored by incretin analog therapy (**Fig. 3D-F**), although incretin analog therapy restored glucose tolerance and insulin sensitivity (**Fig. 1G-H**). These observations suggest that other insulin-sensitive organs compensate for the reduced capacity of adipose tissue to maintain glucose homeostasis. It has been shown that GLP-1 inhibits glucagon secretion and thereby hepatic glucose production (8, 34). Thus, semaglutide and tirzepatide might exert a similar inhibition of hepatic glucose production and thereby improve glucose homeostasis, despite of impaired adipose DNL.

We observed a significant increase in food intake in the first week after incretin analog withdrawal (**Fig. 5D**). Interestingly, food intake in incretin therapy withdrawn mice was higher than in ND or HFD-fed control mice (**Fig. 5D**). These data suggest the presence of a feedback mechanism to increase appetite and thereby restore bodyweight loss caused by incretin analog therapy. Although the underlying mechanism is unknown, the appetite-suppressing hormone leptin and leptin responsiveness in hypothalamus – the central appetite controller – might play a role in the increased food intake upon incretin therapy withdrawal. It has been demonstrated that leptin and the GLP-1 receptor-agonist liraglutide synergistically act on the hypothalamus to reduce appetite and body weight (35). Another study also demonstrated that leptin treatment synergizes with the GLP-1 receptor agonist exendin-4 and a dietary intervention to suppress food intake, leading to bodyweight loss (36). Although these previous studies demonstrate the importance of co-agonism of GLP-1 and leptin receptors for weight loss, there is no information about leptin responsiveness after incretin therapy withdrawal. We observed similar levels of leptin mRNA in adipose tissue and serum leptin in mice treated with incretin analogs, compared to treatment naïve ND-fed mice (**Fig. 2E** and **S2C**). Thus, leptin responsiveness in the hypothalamus, rather than circulating leptin levels, may account for the increased food intake upon incretin therapy withdrawal.

There are currently no clear guidelines for follow-up after incretin analog-induced weight loss. Here we show that an energy-restricted dietary intervention following semaglutide or tirzepatide withdrawal minimized bodyweight gain, maintained glucose tolerance and restored adipose tissue function. Interestingly, a dietary intervention after incretin analog withdrawal increased lean mass (**Fig. 5F**), which is known to counteract bodyweight regain and contributes to maintain glucose homeostasis (37, 38). In patients, significant loss of lean mass loss has been described after semaglutide (39) and tirzepatide treatment (40). Thus, our findings propose that a dietary intervention might be a treatment option to preserve lean body mass after weight loss upon incretin analog treatment.

It is challenging for patients with obesity to adhere to a dietary intervention for weight loss, due to compensatory mechanisms including increased appetite, decreased energy expenditure (41), and the presence of an obesogenic memory (5). Incretin analogs suppress food intake and hedonic appetite (42–45). One study showed that rodents treated with tirzepatide preferentially suppressed intake of lipids, without decreasing intake of chow or sucrose (46). Thus, incretin analog therapy may help for patients with obesity to establish a healthy dietary habit during incretin therapy.

Altogether, our findings highlight the importance of a high-quality dietary intervention as a follow-up treatment after incretin analog withdrawal to restore adipose tissue functions with increased lean mass. Thus, in clinical practice, a dietary intervention in combination with incretin therapy may circumvent drug shortage but also overcome the psychological and financial discomfort of chronic incretin analog therapy for patients with obesity, which ultimately improves patients’ quality of life.

## ACKNOWLEDGEMENTS

We thank Alvaro Cortes Calabuig (Genomics Core KU Leuven, Leuven) for help with RNA sequencing analysis and Carla Cadena del Castillo and Femke van Geest for scientific input.

## DISCLAIMERS

N/A.

## AUTHOR CONTRIBUTIONS

J.S., B.D., G.C., B.V.D.S., S.S., and M.S. conceived and designed research; J.S., B.D., J.D.G., F.V.L., I.S, K.M. and M.S. performed experiments; J.S. and M.S. analyzed data; J.S. and M.S. interpreted results of experiments; J.S. prepared figures; J.S. and M.S. drafted manuscript; all authors edited and revised manuscript and approved the final version of manuscript.

---

**Table S1:**
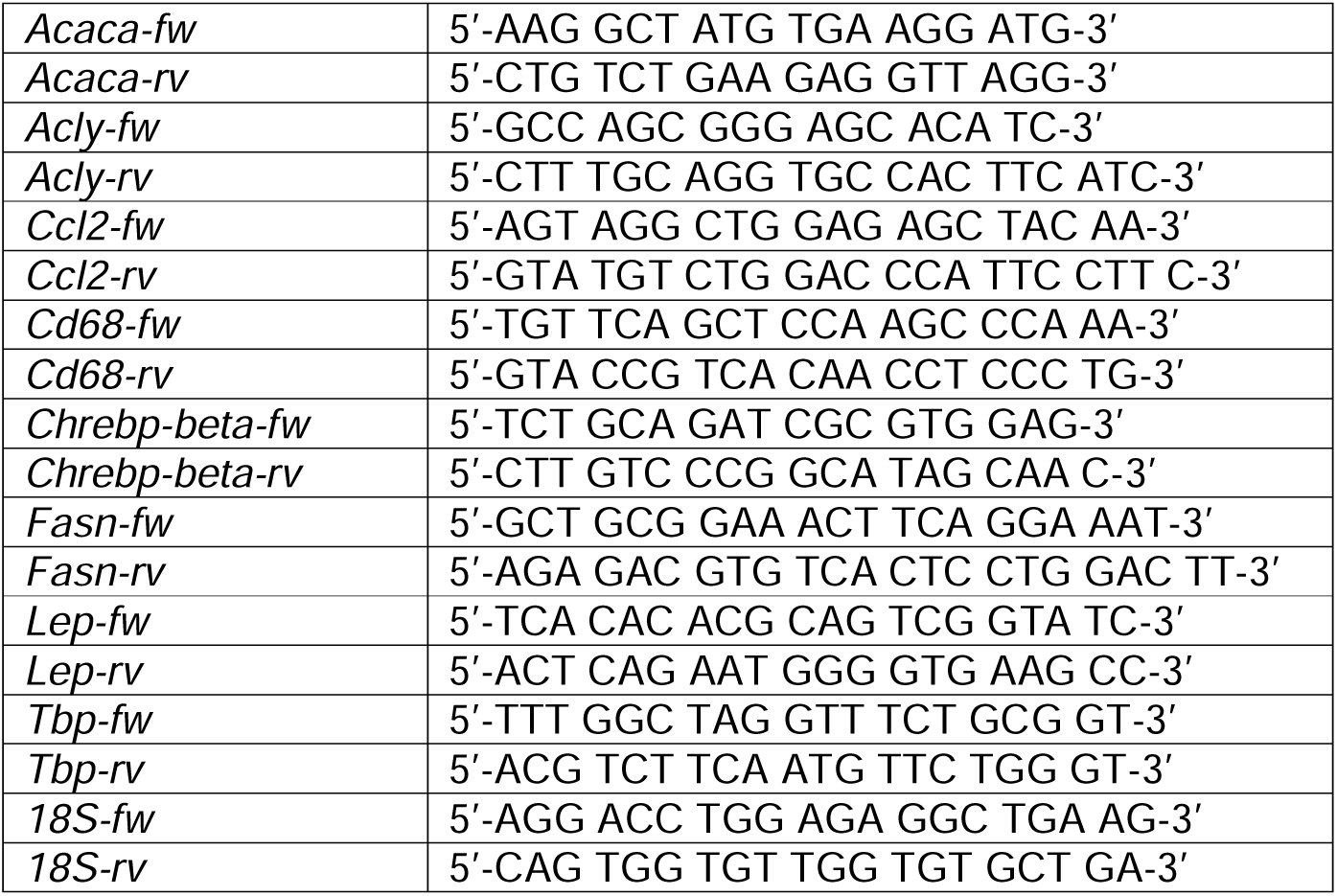
Primers used for RT-qPCR analysis.

