## Supplementary Figures for "A dietary intervention following incretin analog treatment restores adipose tissue functions in diet-induced obese mice"

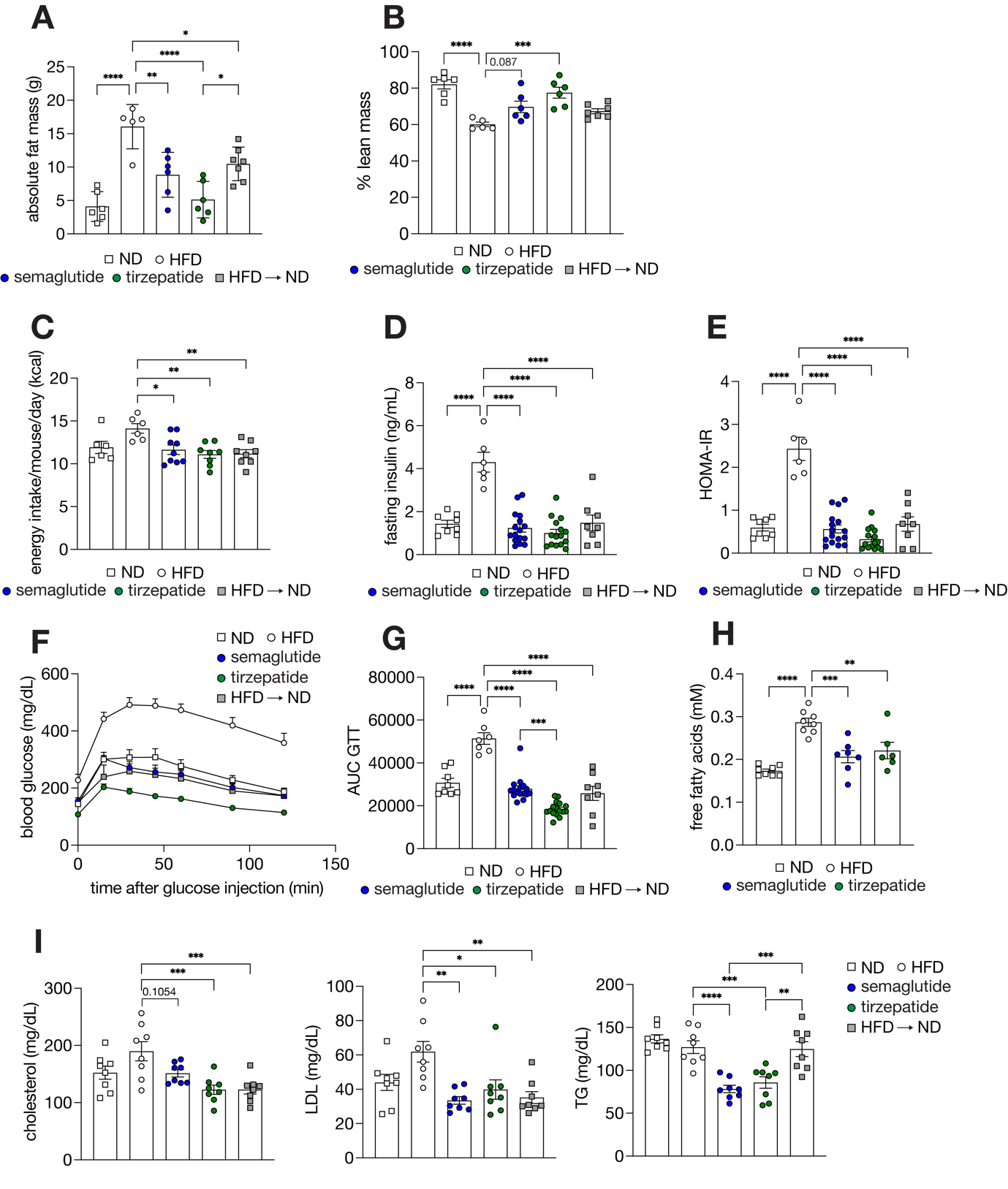

**Figure S1**. **Incretin analog treatment reduces bodyweight and improved glucose tolerance and serum lipid profile. A-B.** Absolute fat mass (A) and relative lean mass (B) after 4 weeks of incretin analog treatment or a dietary intervention. n = 6 (ND, TZP, SEM), 5 (HFD) and 7 (HFD 🡪 ND). **C.** Energy intake was calculated based on food intake (Fig. 1G). N = 6 (ND and HFD), 9 (SEM), and 8 (TZP and HFD 🡪 ND). **D-E**. Fasting insulin (C) and HOMA-IR levels (D) after 4 weeks of incretin analog treatment or a dietary intervention. n = 8 (ND and HFD 🡪 ND), 6 (HFD) and 16 (SEM and TZP) **F-G.** Blood glucose levels during glucose tolerance test (E) and Area under the curve (AUC) (F). 1g/kg glucose was injected, and glucose concentrations were measured at different timepoints. n = 8 (ND and HFD 🡪 ND), 7 (HFD) and 16 (SEM and TZP) **H.** Serum free fatty-acids 8 weeks after incretin analog treatment. n = 8 (ND and HFD), 7 (SEM) and 6 (TZP) **I.** Serum lipid levels (cholesterol, LDL and TG) 4 weeks after incretin analog treatment or a dietary intervention. n = 8. HOMA-IR: homeostatic model assessment of insulin resistance, LDL: low-density lipoprotein, TG: triglycerides *p<0.05, ** p<0.01, ***p<0.001, ****p <0.0001, One-Way ANOVA.

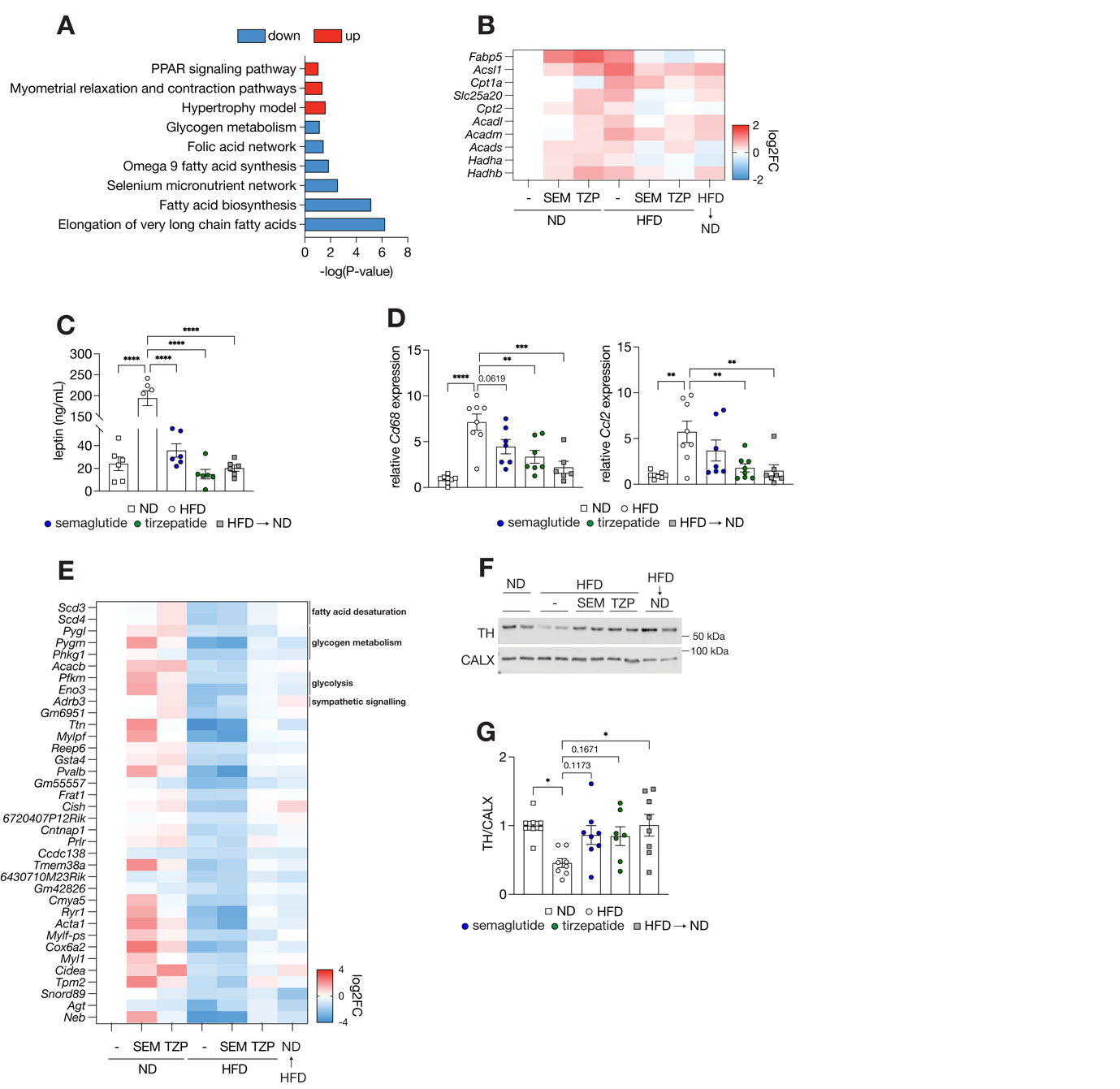
**Figure S2. Incretin analogs restore most of the obesity-mediated deregulated genes in adipose tissue. A.** Pathway analysis of genes with a down- or upregulated expression upon an HFD **B.** Relative expression of genes involved in fatty acid oxidation in sWAT. n = 7 (ND, and ND SEM), n = 8 (HFD, HFD SEM, HFD TZP and HFD 🡪 ND). (**C**) Serum leptin levels 4 weeks after incretin analog treatment or dietary intervention. n = 6 (ND), n = 4 (HFD), n = 6 (SEM, TZP and HFD 🡪 ND) **D.** Relative expression of *Cd68* and *Ccl2* in vWAT. n = 8 **E.** Relative expression of genes that were restored upon TZP or a dietary intervention in sWAT. n = 7 (ND, and ND SEM), n = 8 (HFD, HFD SEM, HFD TZP and HFD 🡪 ND) **F-G.** Immunoblot analysis of tyrosine hydroxylase (TH). CALX serves as a loading control (F) and quantification (G) n = 8. *p<0.05,** p<0.01,***p<0.001, ****p<0.0001 One-Way ANOVA.

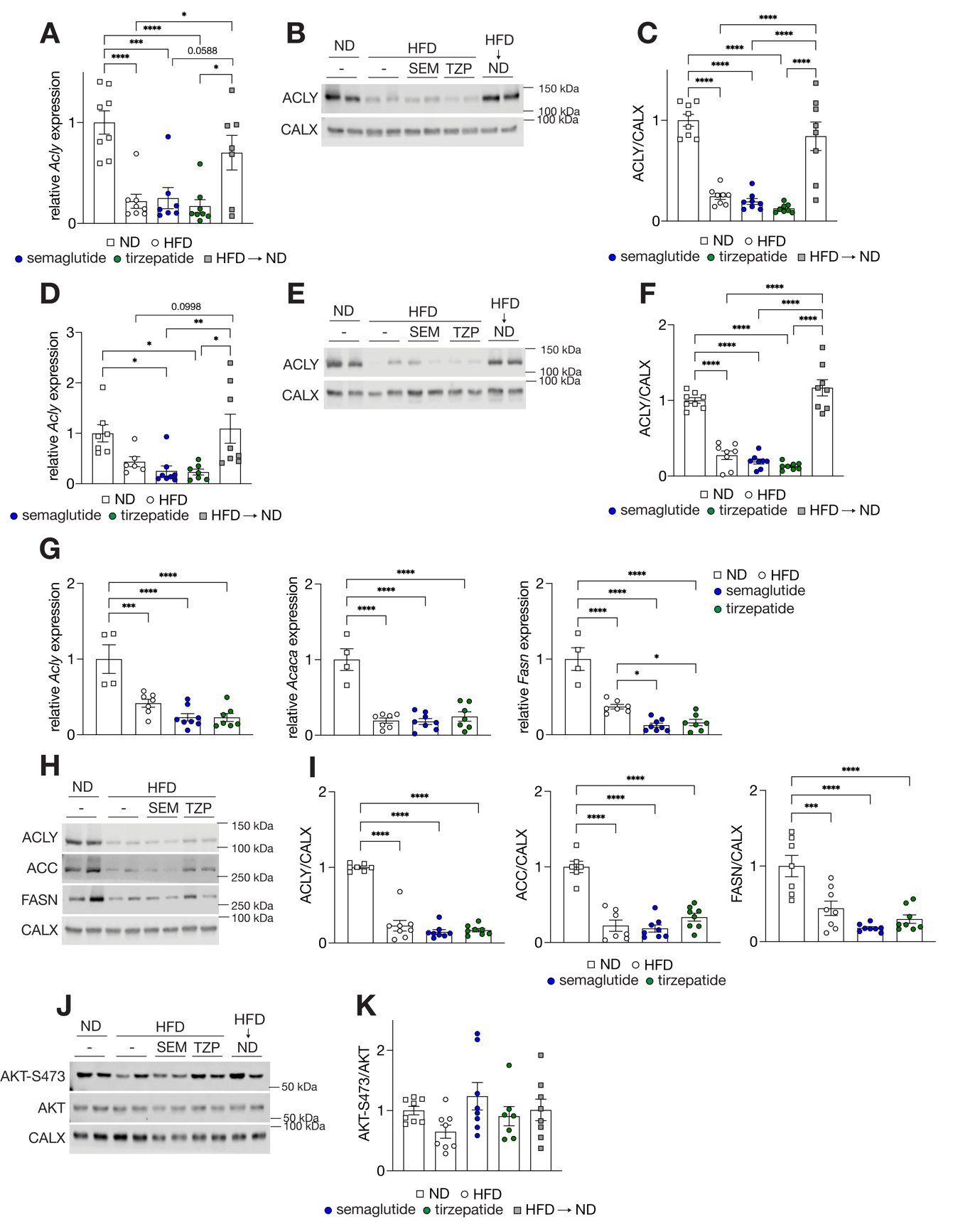

**Figure S3. Incretin analogs does not restore adipose DNL A.** Relative expression of ATP citrate lyase (*Acly*) in vWAT 4 weeks after treatment with incretin analogs or a dietary intervention. n = 8 **B-C.** Immunoblot analysis of ACLY in vWAT 4 weeks after treatment with incretin analogs or a dietary intervention. (B) and quantification (C). n = 8 **D.** Relative expression of Acly in brown adipose tissue 4 weeks after treatment with incretin analogs or a dietary intervention. n = 8 **E-F.** Immunoblot analysis of ACLY in brown adipose tissue 4 weeks after treatment with incretin analogs or a dietary intervention. (E) and quantification. n = 8 (F). **G.** Relative expression of *Acly*, *Acaca*, and *Fasn* after 8 week incretin analog treatment for 8 weeks. n = 4 (ND), n = 7 (HFD, SEM and TZP.) **H-I**. Immunoblot analysis of sWAT ACLY, ACC and FASN in sWAT (H) and quantification (I) after 8 week incretin analog treatment for 8 weeks. n = 7 (ND), n = 8 (HFD, SEM and TZP) **J-K.** Immunoblot analysis of sWAT AKT and AKT phosphorylated at S473 in sWAT. (J) and quantification (K) 4 weeks after treatment with incretin analogs or a dietary intervention. n = 8. *p<0.05,** p<0.01,***p<0.001, ****p<0.0001 One-Way ANOVA.

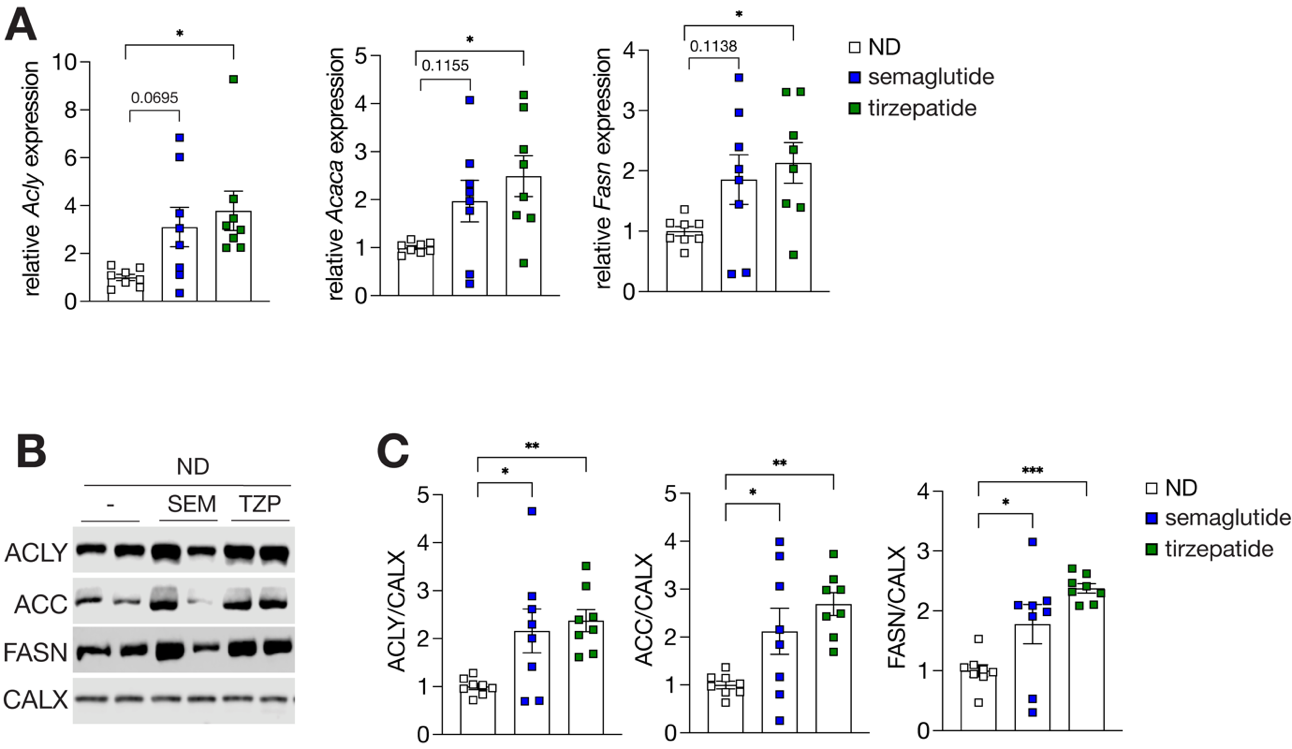

**Figure S4. Incretin analog treatment upregulates adipose DNL gene expression in ND-fed mice. A**. Relative expression of sWAT *Acly*, *Acaca*, and *Fasn* 4 weeks after incretin analog treatment of ND-fed mice **B-C.** Immunoblot analysis of lipogenic enzymes 4 weeks after incretin analog treatment of ND-fed mice (B) and quantification (C). n = 8, ND: normal diet, *p<0.05,**p<0.01, ***p<0.001, ****p<0.0001 , One-Way ANOVA.

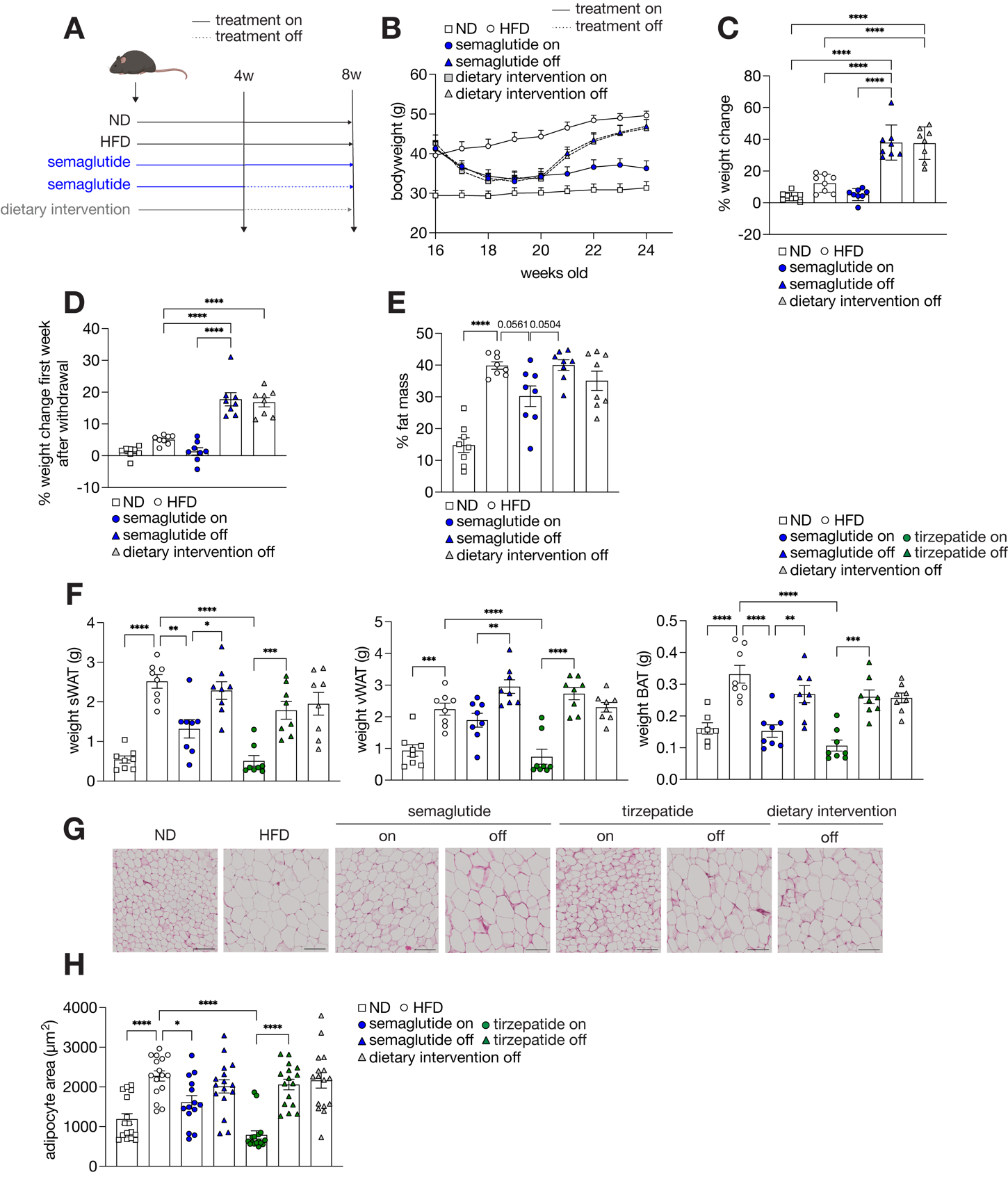
**Figure S5. Incretin analog therapy withdrawal increases bodyweight and adiposity A.** Experimental design of therapy withdrawal **B.** Bodyweight curve duringbefore and after therapy withdrawal or continuous SEM treatment or after withdrawal with dietary switch as control. n = 8. The dotted line indicates therapy withdrawal. ND, HFD control and dietary intervention off groups are representing same samples as in Fig. 4B. **C-D**. Percentage of bodyweight change 4 weeks after therapy withdrawal (C) and after the first week of therapy withdrawal (D). n = 8. (ND, HFD control and dietary intervention off groups are representing same samples as in Fig. 4C-D) **E-F.** Relative fat mass (E) (ND, HFD control and dietary intervention off groups are representing same samples as in Fig. 4E) and adipose depot weight (F) after 4 weeks of therapy withdrawal or continuous incretin analog treatment. n = 8 **G-H**. Hematoxylin and eosin staining of sWAT. bar = 100 µm (G) and quantification (H) Two sections per for each animal were analyzed using Adiposoft-software in ImageJ and the average adipocyte area was calculated for each sectionslice. n = 8. BAT: brown adipose tissue *p<0.05,**p<0.01, ***p<0.001, ****p<0.0001 , One-Way ANOVA

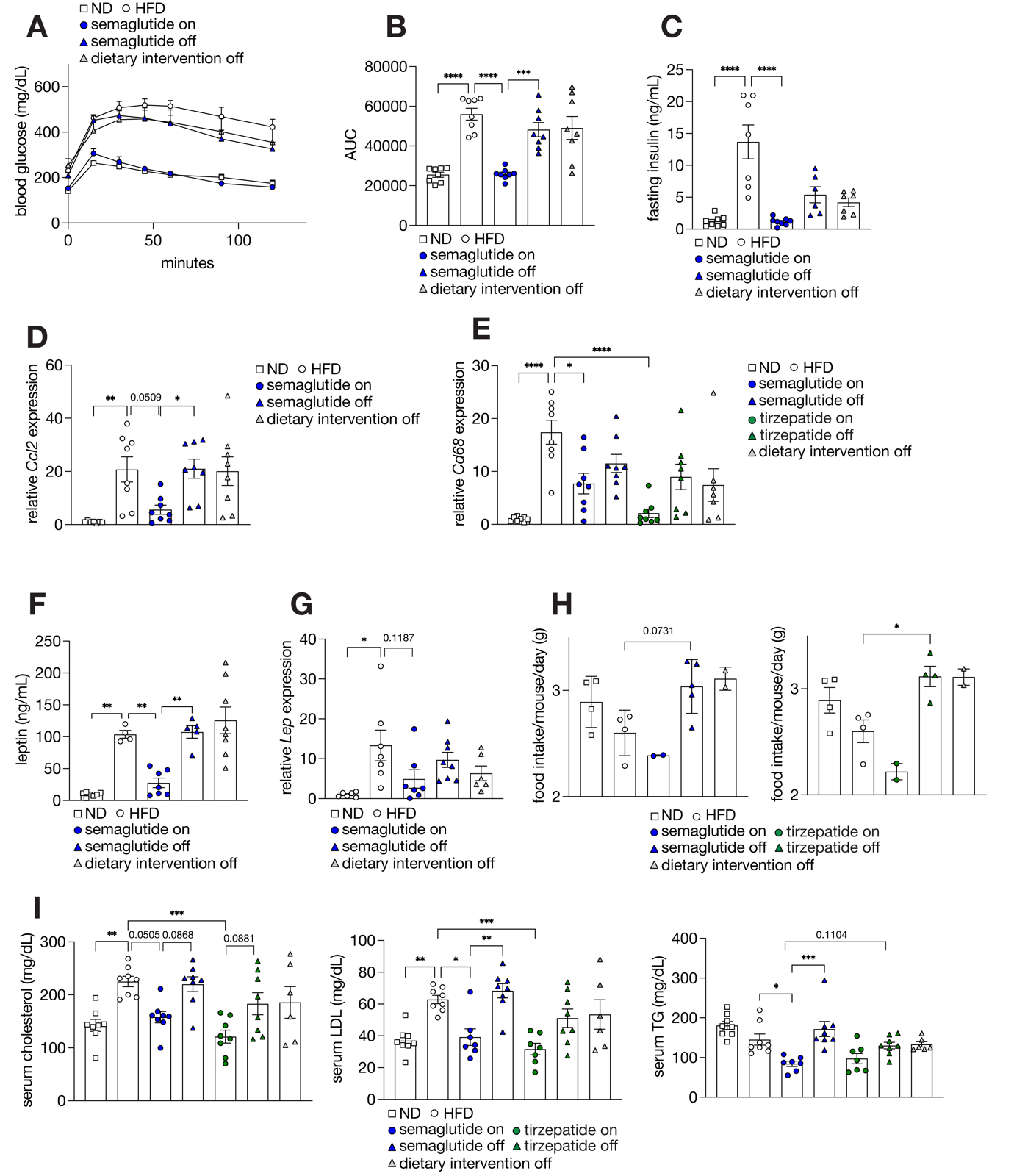
**Figure S6. Incretin analog therapy withdrawal impairs adipose function A-B.** Blood glucose levels during glucose tolerance test 4 weeks after therapy withdrawal or continuous SEM treatment (A) and AUC (B) 1g/kg glucose was injected, and glucose concentrations were measured at different timepoints. n = 8. ND, HFD control and dietary intervention off groups are representing same samples as in Fig. 5F-G **C.** Fasting insulin levels 4 weeks after therapy withdrawal or continuous SEM treatment n = 8 (ND, SEM), 7 (HFD, dietary intervention off) and 6 (SEM off). ND, HFD control and dietary intervention off groups are representing same samples as in Fig. 5H. **D-E.** Relative expression of *Ccl2* (D) and *Cd68* (E) in vWAT. n = 8 **F-G.** Serum leptin levels (F) and relative expression of *Lep* (encoding leptin) (G) in sWAT 4 weeks after therapy withdrawal or continuous SEM treatment. n = 8 (ND, dietary intervention off), 4 (HFD), 6 (SEM) and 5 (SEM off). ND, HFD control and dietary intervention off groups are representing same samples as in Fig. 5J-K. **H.** Food intake 4 weeks after therapy withdrawal or continuous incretin analog treatment. N = 4 (ND, HFD and TZP off), 2 (SEM, TZP and dietary intervention off), 5 (SEM off). ND, HFD control and dietary intervention off groups are representing same samples in both graphs **I.** serum lipid profile (cholesterol, LDL and TG) 4 weeks after therapy withdrawal or continuous incretin analog treatment. n = 8. LDL: low-density lipoprotein, TG: triglycerides, *p<0.05,**p<0.01, ***p<0.001, ****p<0.0001 , One-Way ANOVA

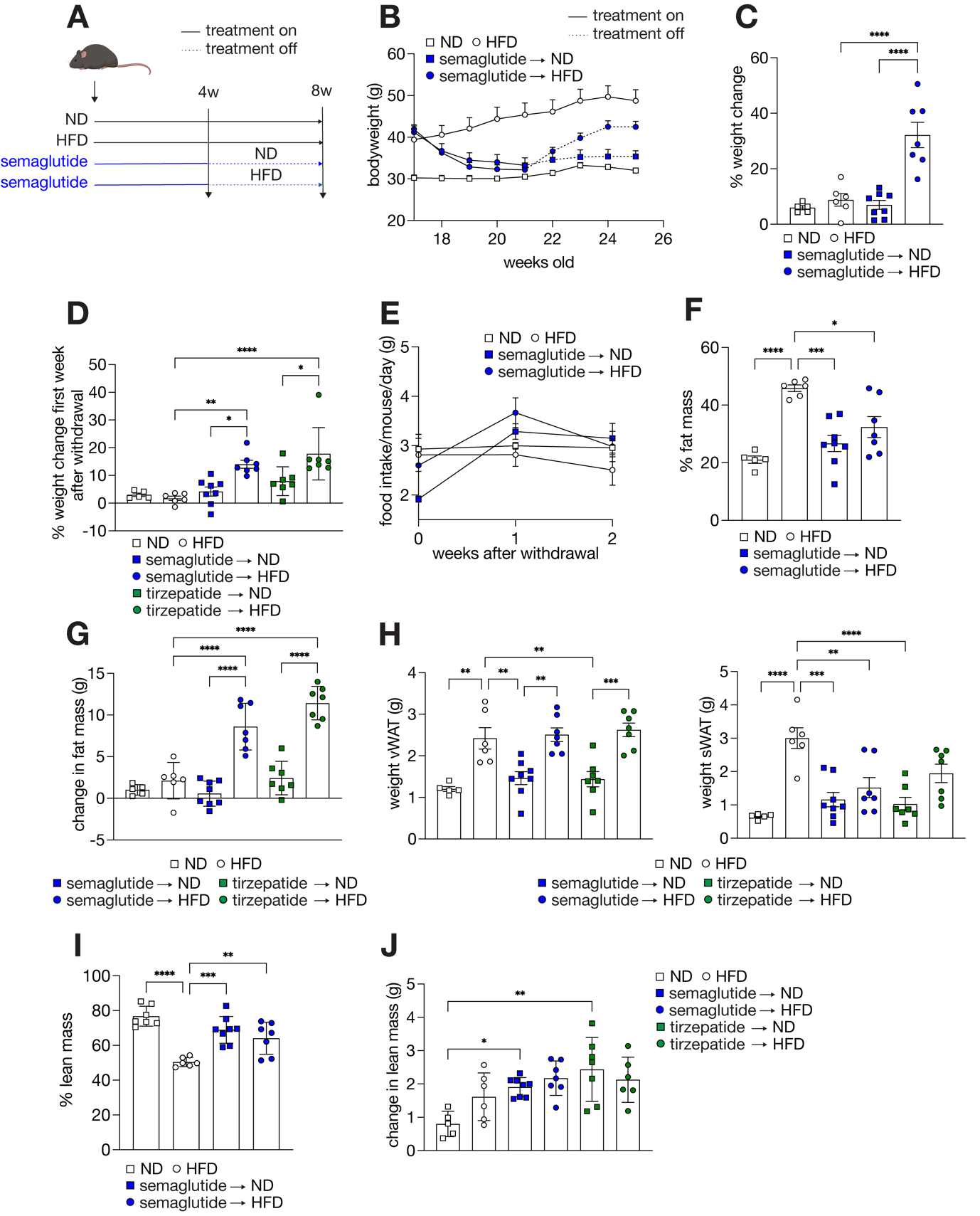

**Figure S7**. **A dietary intervention following incretin analog treatment minimizes bodyweight gain and improves adiposity A.** Experimental design of dietary intervention after incretin withdrawal. **B.** Bodyweight of mice that were maintained on a HFD or switched to a ND for 4 weeks after SEM withdrawal. The dotted line indicates therapy withdrawal. n = 5 (ND), 6 (HFD), 8 (SEM 🡪 ND) and 7 (SEM 🡪 HFD). ND and HFD control groups are representing same samples as in Fig. 5B. **C-D.** Percentage of bodyweight change after 4 weeks (C) and during the first week of incretin analog withdrawal (D) n = 5 (ND), 6 (HFD), 8 (SEM 🡪 ND) and 7 (SEM) 🡪 HFD, TZP 🡪 ND and TZP 🡪 HFD) **E.** Food intake in the first and second week after SEM withdrawal. ND and HFD control groups are representing same samples as in Fig. 5D. **F-G.** Relative fat mass (F) and change in fat mass (ND and HFD control groups are representing same samples as in Fig. 5E) (G) 4 weeks after therapy withdrawal. n = 5 (ND), 6 (HFD), 8 (SEM 🡪 ND) and 7 (SEM) 🡪 HFD). **H.** Weight of vWAT and sWAT. n = 5 (ND), 6 (HFD), 8 (SEM 🡪 ND) and 7 (SEM) 🡪 HFD, TZP 🡪 ND and TZP 🡪 HFD). **I-J.** Relative lean mass (ND and HFD control groups are representing same samples as in Fig. 5F) (I) and change in lean mass (J) 4 weeks after therapy withdrawal. n = 5 (ND), 6 (HFD), 8 (SEM 🡪ND) and 7 (SEM) 🡪 HFD, TZP 🡪 ND and TZP 🡪 HFD). *p<0.05,**p<0.01, ***p<0.001, ****p<0.0001 , One-Way ANOVA

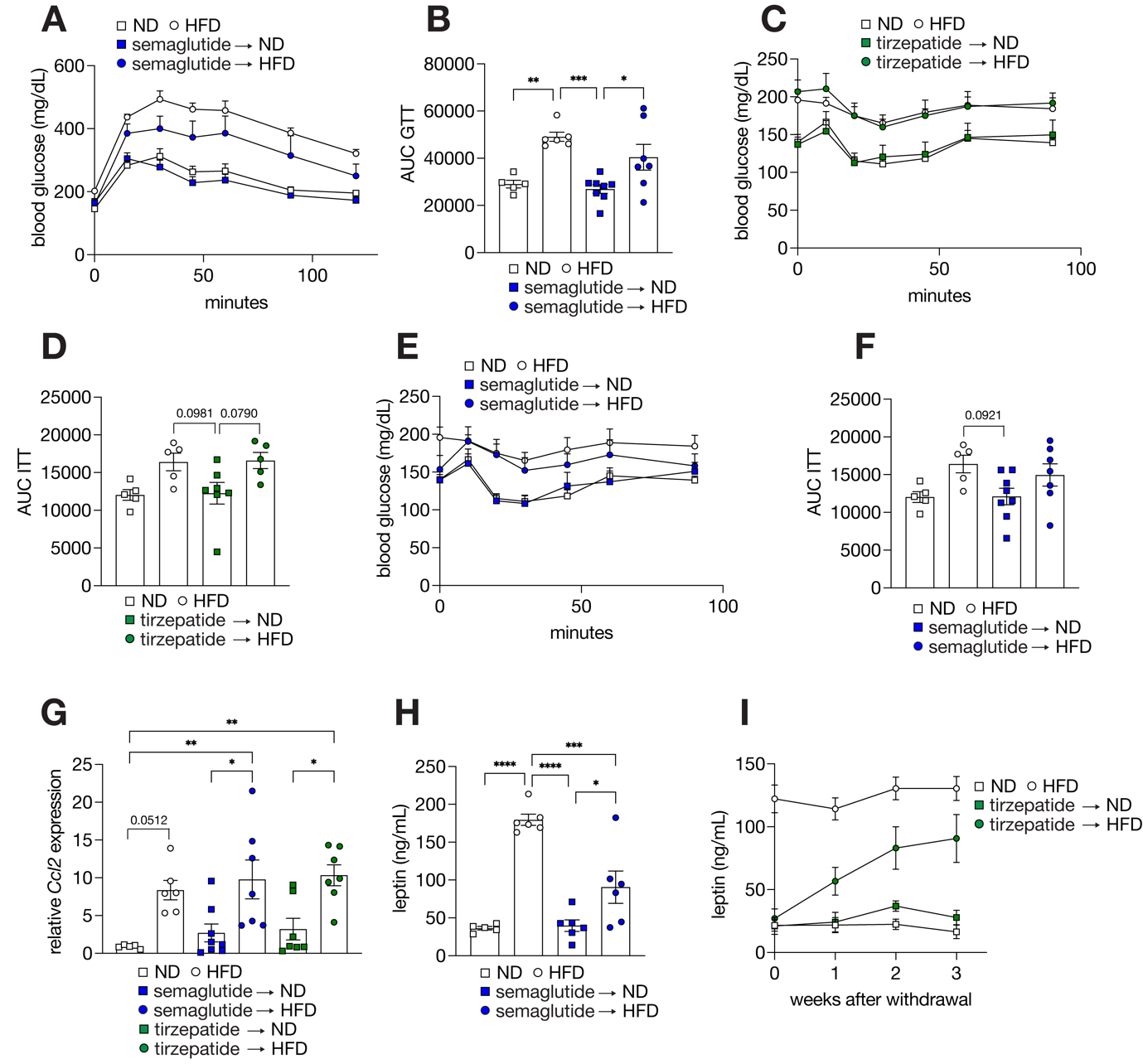

**Figure S8. A dietary intervention following incretin analog treatment maintains glucose homeostasis and adipose tissue function. A-B.** Blood glucose levels during glucose tolerance test 4 weeks after SEM withdrawal (A) and AUC (B). 1g/kg glucose was injected, and glucose concentrations were measured at different timepoints. n = 5 (ND), 6 (HFD), 8 (SEM 🡪 ND) and 7 (SEM 🡪 HFD). ND and HFD control groups are representing same samples as Fig. 5G-H. **C-D.** Blood glucose levels during insulin tolerance test 4 weeks after TZP withdrawal (C) and AUC (D) n = 5 (ND, HFD, TZP 🡪 HFD) and 7 (TZP 🡪 ND) **E-F.** Blood glucose levels during insulin tolerance test 4 weeks after SEM withdrawal (E) and AUC (F). 0.5U/kg insulin was injected, and glucose concentrations were measured at the indicated timepoints. n = 5 (ND, HFD), 8 (SEM 🡪 ND) and 7 (SEM 🡪 HFD). ND and HFD control groups are representing same samples as Fig. S8C-D. **G.** Relative expression of *Ccl2* in vWAT 4 weeks after incretin analog therapy withdrawal. n = 5 (ND), 6 (HFD), 7 (SEM 🡪 ND, SEM 🡪 HFD, TZP 🡪 ND and TZP 🡪 HFD) **H.** Serum leptin levels 4 weeks after SEM withdrawal. n = 5 (ND), 6 (HFD, SEM 🡪 ND, SEM 🡪 HFD). ND and HFD control groups are representing same samples as Fig. 5I. **I.** Increase in serum leptin levels during first and second week after TZP withdrawal. n = 6. *p<0.05,**p<0.01, ***p<0.001, ****p<0.0001 , One-Way ANOVA

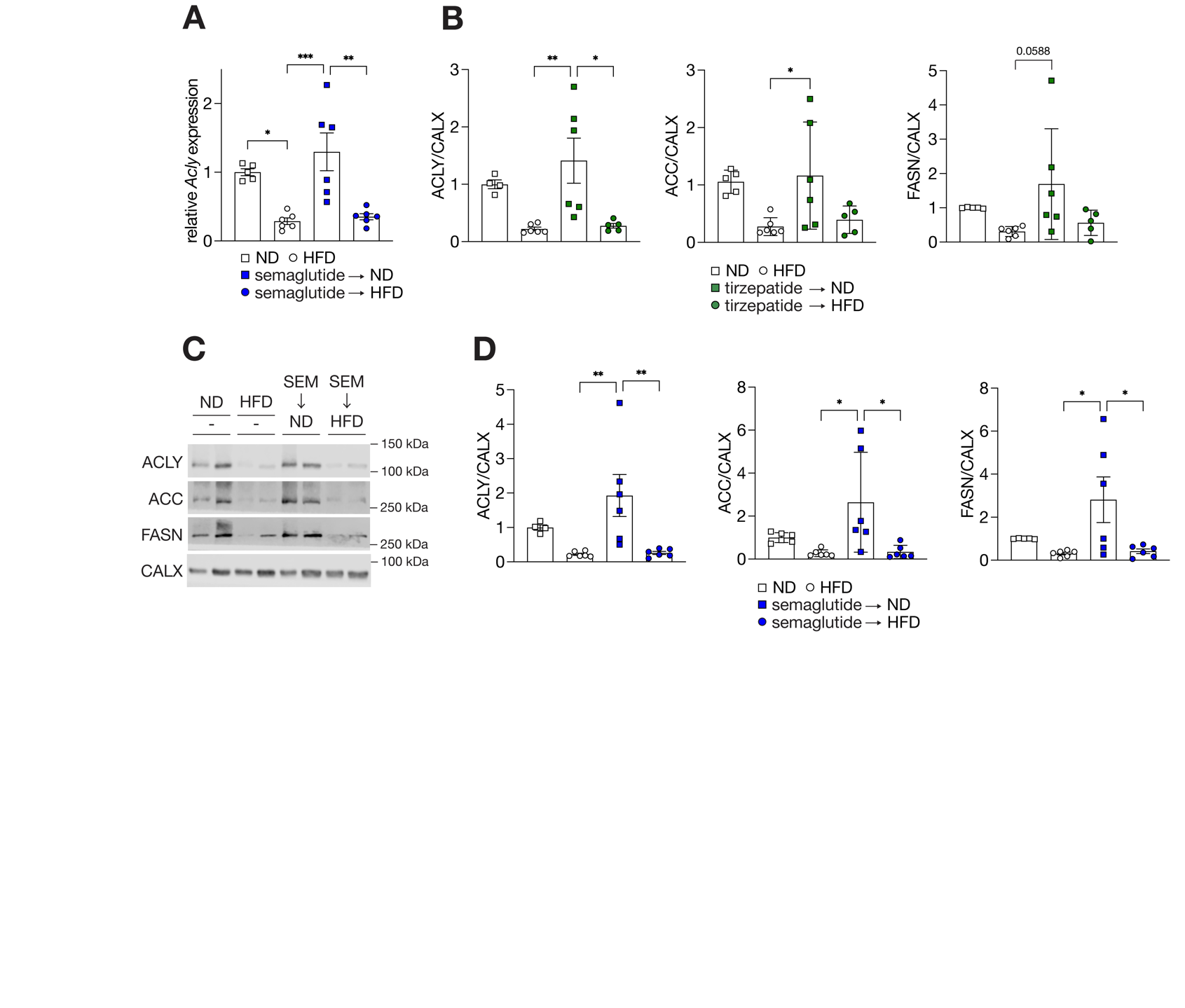
**Figure S9. A dietary intervention following incretin analog treatment restores adipose DNL A.** Relative expression of *Acly* after SEM withdrawal. n = 5 (ND), n = 6 (HFD, SEM 🡪 ND and SEM 🡪 HFD). ND and HFD control groups are representing same samples as in Figure 5J **B.** Quantification of immunoblot in Fig. 5K. n = 5 (ND), 6 (HFD, TZP 🡪 ND, TZP 🡪 HFD) **C-D.** Immunoblot analysis of ACLY, ACC and FASN in sWAT 4 weeks after SEM withdrawal (C) and quantification (D). n = 4 (ND), n = 6 (HFD, SEM 🡪 ND and SEM 🡪 HFD). ND and HFD control groups are representing same samples as in Fig. 5K. *p<0.05,**p<0.01, ***p<0.001, ****p<0.0001, One-Way ANOVA.
